# Broad-scale phenotyping in Arabidopsis reveals varied involvement of RNA interference across diverse plant-microbe interactions

**DOI:** 10.1101/2024.06.25.600702

**Authors:** Alessa Ruf, Hannah Thieron, Sabrine Nasfi, Bernhard Lederer, Sebastian Fricke, Trusha Adeshara, Johannes Postma, Patrick Blumenkamp, Seomun Kwon, Karina Brinkrolf, Michael Feldbrügge, Alexander Goesmann, Julia Kehr, Jens Steinbrenner, Ena Šečić, Vera Göhre, Arne Weiberg, Karl-Heinz Kogel, Ralph Panstruga, Silke Robatzek, the exRNA consortium

**Author notes:** These authors contributed equally to the study.

## Abstract

RNA interference (RNAi) is a crucial mechanism that can contribute to immunity against infectious microbes through the action of DICER-LIKE (DCL) and ARGONAUTE (AGO) proteins. In the case of the fungal pathogen *Botrytis cinerea* and the oomycete *Hyaloperonospora arabidopsidis*, plant DCL and AGO proteins have proven roles as negative regulators of immunity, suggesting functional specialization of these proteins. To address this aspect in a broader taxonomic context, we characterized the colonization pattern of an informative set of *DCL* and *AGO* loss-of-function mutants in *Arabidopsis thaliana* upon infection with a panel of pathogenic microbes with different lifestyles, and a fungal mutualist. Our results revealed that AGO1 and AGO4 function as positive regulators of immunity to a bacterial and a fungal pathogen, respectively. Additionally, AGO2 and AGO10 positively modulated the colonization by a fungal mutualist. Therefore, analysing the role of RNAi across a broader range of plant-microbe interactions has identified previously unknown functions for AGO proteins. For some pathogen interactions, however, all tested mutants exhibited wild type-like infection phenotypes, suggesting that the roles of AGO and DCL proteins in these interactions may be more complex to elucidate.

## INTRODUCTION

RNA interference (RNAi) is a conserved mechanism that regulates gene expression via small (s)RNAs (Huang et al., 2019; Tang et al., 2022). These regulatory molecules ranging from 19– 24 nucleotides (nt) are mainly classified into micro (mi)RNAs (20-22 nt duplexes) and small interfering (si)RNAs (21-24 nt duplexes), which is based on their biogenesis pathways (Axtell, 2013). miRNAs are produced from imperfect stem-loop structures of non-coding (nc)RNAs and typically mediate post-transcriptional gene silencing (PTGS) of sequence-complementary mRNA targets. By contrast, siRNAs originate from double-stranded (ds)RNAs and either direct PTGS of the mRNA target or guide RNA-directed DNA methylation (RdDM) for transcriptional gene silencing (TGS) (Martín-Merchán et al., 2023).

In the interaction with infectious agents, host sRNAs target foreign genes to mediate defence, e.g. against viruses (Obbard et al., 2008; Zhan & Meyers, 2023). Host sRNAs also fine-tune the expression of host immune-responsive genes, thereby orchestrating the outcome of infection against various pathogens (Šečić, Kogel, et al., 2021). For example, in the genetic model *Arabidopsis thaliana*, miRNA393 contributes to resistance against *Pseudomonas syringae pv tomato* strain DC3000 (*Pto* DC3000) by regulating pattern-triggered immunity (PTI) through the suppression of auxin signalling (Navarro et al., 2006). During seedling development, miR172 inhibits the expression of FLAGELLIN SENSING2 (FLS2), a well-studied pattern recognition receptor (PRR) that confers PTI against flagellated bacteria (Zou et al., 2018). This suggests a role of miR172 in coordinating plant immunity and development.

The core mechanism of RNAi involves the production of dsRNAs, which are processed by DICER-LIKE (DCL) proteins into sRNA duplexes. These sRNAs are then methylated by HUA ENHANCER1 (HEN1), and subsequently loaded into RNA-induced silencing complexes (RISCs) (Iwakawa & Tomari, 2022; Martín-Merchán et al., 2023). ARGONAUTE (AGO) proteins are the main components of RISCs and recruit single-stranded sRNAs for pairing with sequence-complementary RNA and DNA targets (Fang & Qi, 2016). To amplify RNAi, RNA-dependent RNA polymerases use single-stranded sRNA for the generation of long dsRNAs, subjected to processing by DCL2/DCL4 to produce secondary phased siRNAs (Curaba & Chen, 2008; Martín-Merchán et al., 2023).

Compared to mammals, plants have an enlarged repertoire of RNAi machinery components. *A. thaliana* encodes four DCL proteins and each generates a specifically sized sRNA (Martín-Merchán et al., 2023). DCL1 produces 21/22 nt miRNAs involved in PTGS. DCL1 is essential since its complete genetic deletion leads to embryonic lethality (Kurihara & Watanabe, 2004). A partial *dcl1* loss-of-function mutant in *A. thaliana* showed enhanced susceptibility to *Pseudomonas syringae* pv. *tomato* (*Pto*) DC3000 and *Botrytis cinerea* infection (Navarro et al., 2006; Weiberg et al., 2013), but also displayed developmental abnormalities. DCL2, DCL3, and DCL4 produce siRNAs of 22 nt, 24 nt, and 21 nt, respectively (Howell et al., 2007). DCL2 generates siRNA from natural *cis*-acting anti-sense transcripts (Borsani et al., 2005) and mediates antiviral immunity (Taochy et al., 2017; Z. Wang et al., 2018). DCL3 is involved in RdDM pathways, processing dsRNAs generated by RDR2 into siRNAs (Coursey et al., 2018). DCL4 produces *trans*-acting siRNAs (Gasciolli et al., 2005), mediating defence against RNA viruses (Azevedo et al., 2010; Bouché et al., 2006; Deleris et al., 2006). Furthermore, DCL4 is also important in anti-fungal defence, since *dcl4* mutants are more susceptible to infections with the vascular fungus *Verticillium dahliae* (Ellendorff et al., 2009).

AGO proteins are widely distributed in eukaryotes, including plants with a very large expansion in monocotyledons, and are also found in prokaryotes (Bobadilla Ugarte et al., 2023). Eukaryotic AGOs typically contain four conserved domains, the N-terminal (N) domain, the PIWI/Argonaute/Zwille (PAZ) domain, the MID and the P-element induced wimpy testis (PIWI) domains, connected by two linker regions, L1 and L2 (Martín-Merchán et al., 2023). The PAZ and MID domains recognize the 3’ and the 5’ ends of sRNAs, respectively, the N-domain interacts with other proteins of the RNAi machinery, and the PIWI domain exhibits RNase catalytic activity for RNA cleavage (Nakanishi, 2024).

Ten AGO proteins have been identified in *A. thaliana* that can be classified into three clades: i) AGO1/5/10 (clade I), ii) AGO2/3/7 (clade II), and iii) AGO4/6/8/9 (clade III) (Martín-Merchán et al., 2023). They feature different subcellular localization patterns and preferences for sRNA binding. Clade I AGO proteins exhibit a predominantly cytoplasmic steady state localization and preferentially bind 21–22 nt siRNAs and miRNAs, which is consistent with their main function in PTGS (Fang & Qi, 2016). AGO1 is also involved in the biogenesis of *trans*-acting siRNA (tasiRNA) and phased secondary RNAs (phasiRNA) (Fang & Qi, 2016). AGO proteins of clade II have likewise a preference for binding 21–22 nt si/miRNAs but exhibit both cytoplasmic and nuclear localizations and play more diverse roles (Martín-Merchán et al., 2023). Functioning mainly in TGS, clade III AGO proteins localize to the nucleus and bind 24 nt sRNAs (Martín-Merchán et al., 2023).

The expression patterns of *AGO* genes does not seem to correlate with their clade assignment and function. The members of clade I and clade III, *AGO1* and *AGO4,* are ubiquitously expressed across tissues and during various developmental stages of *A. thaliana* (Jullien et al., 2022). *AGO5*, *AGO6, AGO7, AGO9*, and *AGO10,* members across all clades, exhibit specific expression patterns with spatial and temporal regulation during plant development (Mallory & Vaucheret, 2010; Wook et al., 2011). The expression of *AGO2* and *AGO3* is induced in response to diverse abiotic and biotic stresses (Martín-Merchán et al., 2023). For example, *AGO2* expression is upregulated during *Pto* DC3000 infection (Zhang et al., 2011). *AGO8* appears to be expressed very lowly and was hypothesized to be a pseudogene, but accumulates in premeiotic ovules of *ago4/ago9* mutants (Hernández-Lagana et al., 2016; Mallory & Vaucheret, 2010).

In addition to the important roles of AGO1 in antiviral defence (Huang et al., 2019; Tang et al., 2022), several studies have examined the roles of AGO proteins in plant immunity against eukaryotic and prokaryotic microbes. For example, specific partial loss-of-function mutants in *AGO1* are compromised in microbe-associated molecular pattern (MAMP)-induced immunity against *Pto* DC3000 (Li et al., 2010). Since infection with the fungal pathogen *Sclerotinia sclerotiorum* showed more severe necrotic disease symptoms in *ago1* mutants (Cao et al., 2020), the study suggests that AGO1 is a positive regulator of PTI and enhances resistance against *S. sclerotiorum*. However, AGO1 has also been described to negatively regulate plant immunity against the fungal pathogens *B. cinerea, V. dahliae, Verticillium longisporum* and *Botryosphaeria dothidea*, and the oomycete *H. arabidopsidis* (Dunker et al., 2020; Ellendorff et al., 2009; Shen et al., 2014; Weiberg et al., 2013; Yu et al., 2017). Yet, AGO1 had no detectable role in the outcome of infection with the fungal and oomycete pathogens *Erysiphe cruciferarum* and *Albugo laibachii*, respectively (Dunker et al., 2020). Of the other clades, Arabidopsis *ago2* mutants are more susceptible to infection by *V. dahliae*, *S. sclerotiorum* and species of the oomycete pathogen *Phytophthora* (Cao et al., 2020; Ellendorff et al., 2009; Guo et al., 2018). Furthermore, AGO4 contributes to resistance to *Pto* DC3000 and is required for both local and *Trichoderma*-induced systemic immunity against *B. cinerea* (Agorio & Vera, 2007; López et al., 2011; Rebolledo-Prudencio et al., 2022).

AGO proteins act together with their loaded sRNAs within the RISC complex, suggesting that the role of AGO proteins as positive or negative regulators of immunity in *A. thaliana* likely depends on the specific sRNAs they are bound to. Beyond the evolution of pathogen-derived molecular suppressors that interfere with host RNAi (Hou et al., 2019; Navarro et al., 2006), infectious microbes can hijack host AGO1 and incorporate microbe-derived sRNAs to facilitate infection. This cross-kingdom (ck)RNAi has been demonstrated for the interaction of *A. thaliana* with the taxonomically diverse pathogens *B. cinerea* and *H. arabidopsidis* (Dunker et al., 2020; Weiberg et al., 2013). In both cases, it is mediated by fungal- or oomycete-derived sRNAs, respectively, which are loaded into host AGO1 and thereby interfere with host RNAi pathways.

In *B. cinerea*, a majority of cross-kingdom sRNAs are derived from Ty3-type retrotransposons that contribute to the pathogenicity of this fungus (Porquier et al., 2021). Consistently, the *B. cinerea rdr1 and dcl1/dcl2* mutants were less virulent on both *A. thaliana* and *Solanum lycopersicum* hosts, since the production of sRNAs was nearly abolished in these fungal mutants (Cheng et al., 2023; Weiberg et al., 2013). Yet, genetic deletion of both *DCL* genes in a *ku70* deletion mutant background or deletion of a genetic region of a multi-copy retrotransposon encoding 10% of the total sRNAs in another *B. cinerea* genotype did not compromise virulence on *A. thaliana*, *S. lycopersicum*, *Nicotiana benthamiana* and *Phaseolus vulgaris* (Qin et al., 2023). Since plants also send sRNAs into *B. cinerea* (Cai et al., 2018), ckRNAi occurs in both directions of the interacting organisms.

In *A. thaliana*, DCL and AGO proteins have proven roles as negative regulators in resistance against both the ascomycete pathogen *B. cinerea* and the oomycete *H. arabidopsidis*, suggesting specialization of these proteins in plant immunity. However, since different *B. cinerea* genotypes exhibited varied infection phenotypes (Qin et al., 2023; Weiberg et al., 2013), the contribution of RNAi to the outcome of microbial infections tends to be more complex and possibly species- or even pathotype-dependent. Therefore, it cannot always be assumed with certainty that plant mutants in the RNAi pathway exhibit phenotypes at each time point when infected with any microbe. To address this aspect in a broader taxonomic context, we characterized the expression patterns and loss-of function mutant phenotypes of an informative set of *DCL* and *AGO* genes upon infection with a panel of pathogenic filamentous microbes and bacteria, each with different lifestyles, including mutualistic colonization. We reproduced some previously investigated phenotypes and uncovered new roles for AGO1, AGO2, AGO4, and AGO10 in certain microbial interactions. However, not all tested interactions revealed obvious phenotypic changes, suggesting functional redundancies within the host *AGO* and *DCL* genes, along with microbial virulence mediated by effectors other than sRNAs.

## RESULTS

We selected a set of *A. thaliana* genes and their corresponding mutants that are informative for the siRNA pathway. These include *DCL2*, *DCL3*, *DCL4* and the triple *dcl2/3/4* mutant and members of the three AGO clades (*AGO1*, *AGO10*, the *ago1-27*, *ago1-46* and *ago10-1* mutants (clade I), *AGO2* and the *ago2-1* mutant (clade II), *AGO4* and the *ago4-2* mutant (clade III) (Supplemental Table S1). Exploring publicly available transcriptome data of *A. thaliana* elicited with microbe-associated molecular patterns (MAMPs) from fungi (ch8, nlp20), oomycete (nlp20) and bacteria (flg22, elf16, LPS, nlp20) (Bjornson et al., 2021), we noted that all tested *AGO* but not the selected *DCL* genes were responsive to the immune stimuli (Supplemental Figure S1). Of the *AGO* genes, *AGO2* showed upregulation in response to all MAMPs, while *AGO1*, *AGO4* and *AGO10* were downregulated in response to bacterial MAMPs. This suggests that *AGO* genes across the three clades could be involved in PTI regulation. The strong MAMP-induced expression of *AGO2* is consistent with its documented role in immunity against *S. sclerotiorum* fungi and anti-bacterial immunity against *Pto* DC3000 and its AvrRpt2-avirulent derivative (Cao et al., 2020; Zhang et al., 2011).

Since *AGO* gene expression was responsive to MAMPs derived from different microbial taxa, we selected a panel of pathogenic fungi (*Thecaphora thlaspeos*, *E. cruciferarum*, *V. longisporum*), a symbiotic fungus (*Serendipita indica*), and bacterial pathogens (*Pto* DC3000, *Xanthomonas campestris* pv. *campestris*, *Xylella fastidiosa* subsp. *fastidiosa*) to study the *DCL* and *AGO* expression profiles as well as the infection phenotypes of corresponding mutants in *A. thaliana* (Supplemental Table S2). We also included an oomycete pathogen (*H. arabidopsidis*), given that AGO1-dependent ckRNAi has been demonstrated to play a role in its infection outcome in *A. thaliana* (Dunker et al., 2020). The selected microbes also differ in their lifestyles, with *H. arabidopsidis* and *Pto* DC3000 infecting leaf mesophyll tissue, *E. cruciferarum* invading leaf epidermal cells, *S. indica* colonizing roots, and *V. longisporum*, *X. campestris pv. campestris* and *X. fastidiosa* infecting the plant xylem, as well as *T. thlaspeos* growing systemically along the vasculature in both roots and aerial tissues (Supplemental Table S2). Appreciating the diverse lifestyles, we performed the infection experiments tailored to the type of plant-microbe interaction and according to well established protocols, yet mainly at the whole plant/organ scale with *in vitro* and soil-grown plants. Gene expression was analysed at different time points of early, middle, and late infection/colonization stages depending on the interacting microbe. The infection/colonization success was measured as the ability to invade host cells (number of penetration events), as microbial biomass (number of hyphae or microbial DNA/RNA), or as the capacity of the microbe to multiply within host tissue (number of colony-forming units (cfu)). To minimize the putative effect of seed batches, we used an age-matched seed collection of *A. thaliana* Col-0 and the selected *dcl2/3/4* and *ago* mutants for our experiments.

### AGO1 is a regulator of immunity against some but not all filamentous pathogens

We first tested our collection of plant lines and investigated the role of *DCL* and AGO proteins in the interaction with *H. arabidopsidis*. Expression of *DCL2*, *DCL4,* and *AGO2* was upregulated at middle (4 days post inoculation (dpi)) and late (6 dpi) stages of *H. arabidopsidis* infection, while *AGO4* was downregulated at these time points (Figure 1A). This is in agreement with the changes in the expression of *AGO2* and *AGO4* in response to the oomycete MAMP nlp20 (Supplemental Figure 1). No drastic changes in gene expression were observed for *AGO1* and *AGO10* (Figure 1A). In the infection experiments, *ago1-27* and *ago1-46* mutants displayed enhanced resistance to *H. arabidopsidis* at 6 dpi (Figure 1B). No altered infection was observed in *ago2-1* and *ago4-2* mutants. This outcome is consistent with a previous report that showed evidence for the loading of pathogen-derived sRNAs into *A. thaliana* AGO1, resulting in ckRNAi to support infection (Dunker et al., 2020). Collectively, the data suggests a specific role for AGO1 in the interaction with the oomycete pathogen.

**Figure 1.**
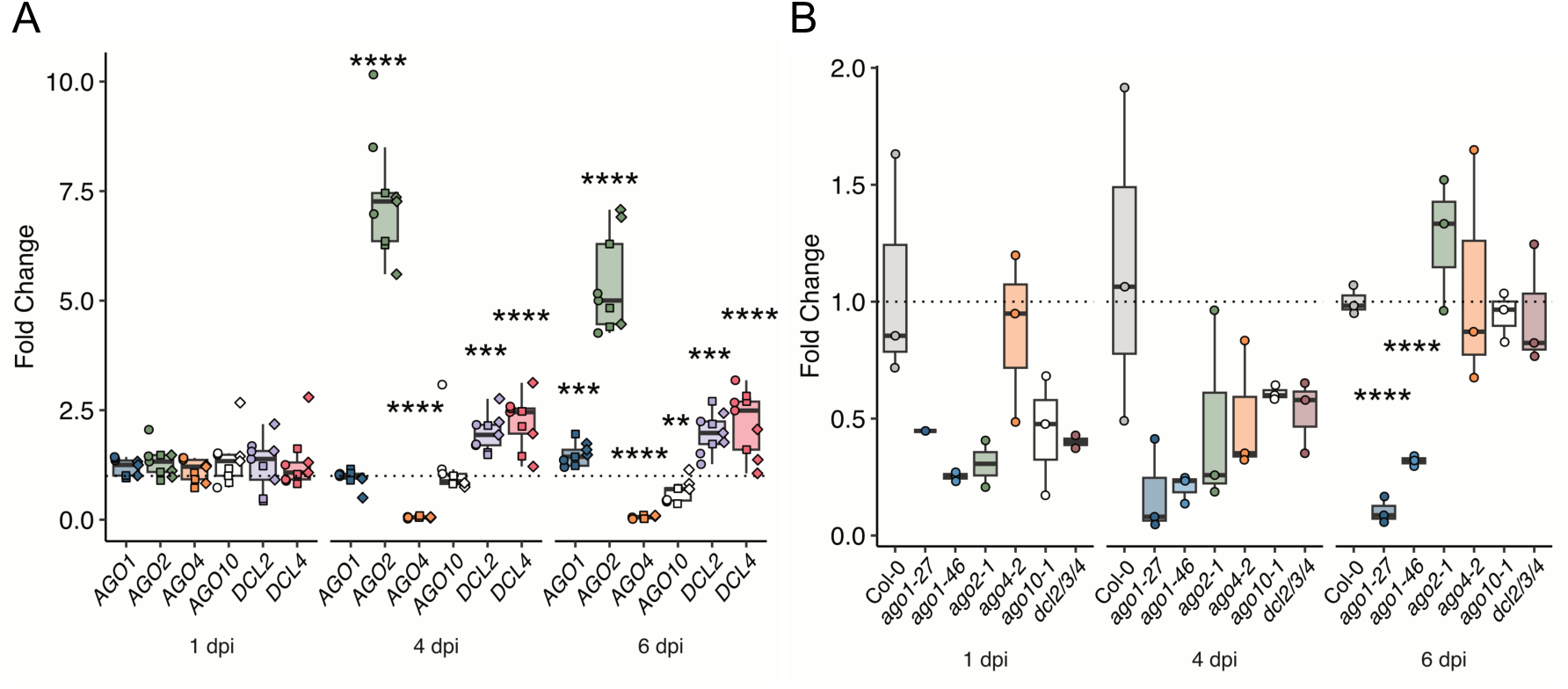
*DCL* and *AGO* gene expression patterns (A) and colonization in respective mutants (B) upon infection with *H. arabidopsidis* isolate Noco 2. **(A)** Samples were collected at 1 dpi (days post inoculation), 4 dpi, and 6 dpi. The RNA levels are relative to mock and normalized against *CDKA*. The results of three independent biological replicates are depicted. **(B)** Pathogen load on *ago* and *dcl* mutants was assessed by measuring relative *H. arabidopsidis* gDNA quantities with RT-qPCR at 1 dpi, 4 dpi, and 6 dpi. The result of one independent biological replicate is depicted. Error bars show standard deviation. Statistical significance was assessed by two-sided Welch’s t-test (α = 0.05, p-values * < .05, ** < .01, *** < 0.001, **** < 0.0001). Symbols indicate number of independent biological replicate. Circle = first, square = second, diamond = third. The dashed line indicates a fold change = 1.

AGO1 also negatively regulates immunity against fungal pathogens including *B. cinerea* and *V. longisporum* but not *E. cruciferarum* (Dunker et al., 2020; Shen et al., 2014; Weiberg et al., 2013). Consistent with the fact that AGO1 negatively regulates immunity against *V. longisporum* (Shen et al., 2014), *AGO1* expression was downregulated in the course of infection with this fungal pathogen (Supplemental Figure S2). By contrast, *DCL3* and *DCL4* were upregulated by *V. longisporum*, suggesting a varied response to this pathogen.

Next, we explored the selected *DCL* and *AGO* genes for their expression profiles in response to challenge with *E. cruciferarum*. We found reduced *AGO1*, *AGO4,* and *AGO10* expression and upregulation of the tested *DCL* genes across the time course (Figure 2A). Fungal entry rates were slightly yet in statistically significant manner increased in *ago1-27*, but no differences were observed in any of the other tested mutants including the allelic *ago1-46* mutant (Figure 2B). This outcome confirms previous data on unaltered *E. cruciferarum* infection in *ago1-46* but contrasts the reported wild type-like phenotype in *ago1-27* (Dunker et al., 2020). This difference could result from the previous study evaluating leaf necrosis while here fungal penetration was scored. Furthermore, a clear regulation of *E. cruciferarum* penetration success by AGO1 cannot be established, since the two *ago1* mutants exhibited different infection phenotypes to this fungus (Figure 2B). We also investigated the role of AGO1 during infection with *T. thlaspeos* and observed wild type-like colonization in *ago1-27* mutants (Figure 3). By contrast, *ago4-2* mutants showed significantly enhanced susceptibility, thereby revealing a previously unknown role for AGO4 as a positive regulator of immunity to this smut fungus.

**Figure 2:**
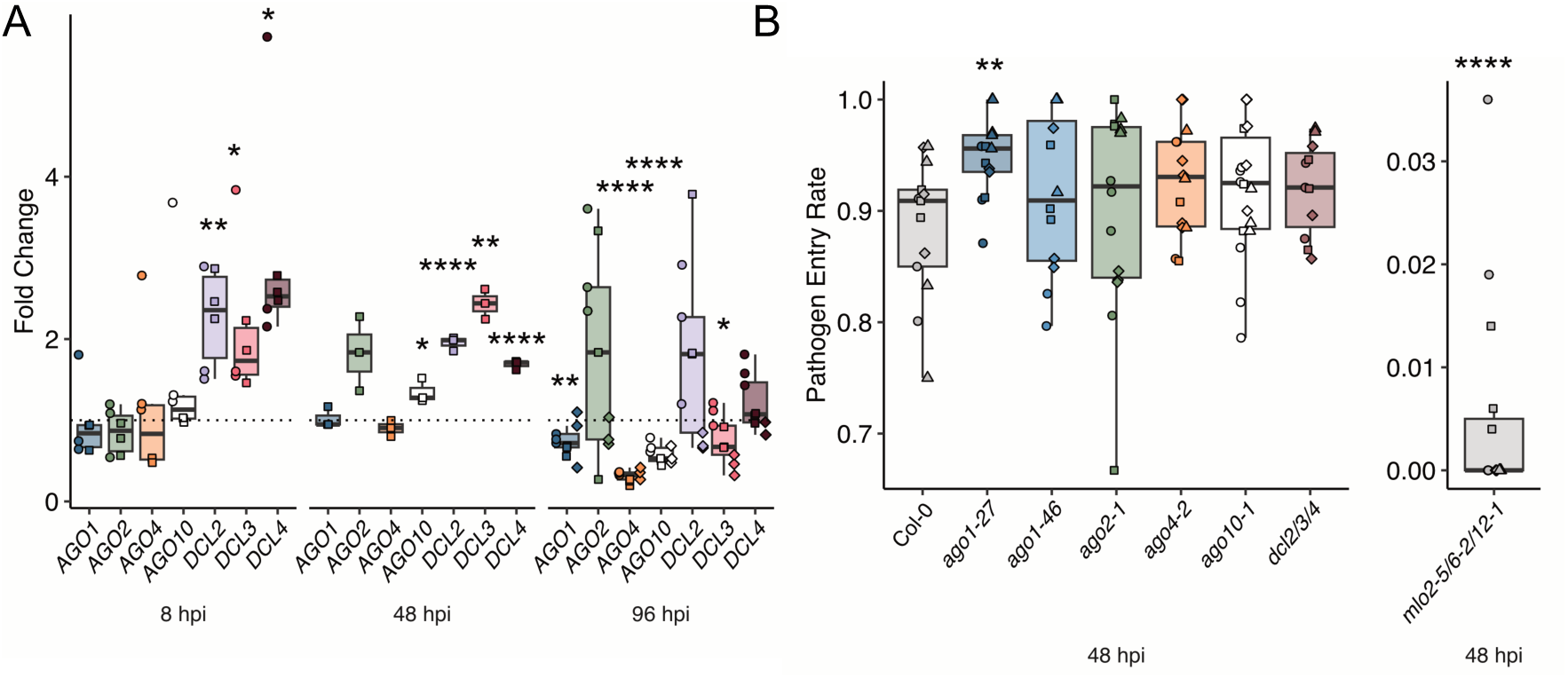
*DCL* and *AGO* gene expression patterns (A) and colonization in respective mutants (B) upon *E. cruciferarum* infection in a time course experiment. **(A)** Samples were collected at 8 hpi (hours post inoculation), 48 hpi, and 96 hpi. The RNA levels are relative to mock and normalized against *CDKA*. The results of three independent biological replicates are depicted. **(B)** Infection success on *ago* and *dcl* mutants was assessed by determining *E. cruciferarum* host cell entry rates at 48 hpi. The results of four independent biological replicates are depicted. Error bars show standard deviation. Statistical significance was assessed by two-sided Welch’s t-test (α = 0.05, p-values * < .05, ** < .01, *** < 0.001, **** < 0.0001). Symbols indicate number of independent biological replicate. Circle = first, square = second, diamond = third, triangle = fourth. The dashed line indicates a fold change = 1.

**Figure 3:**
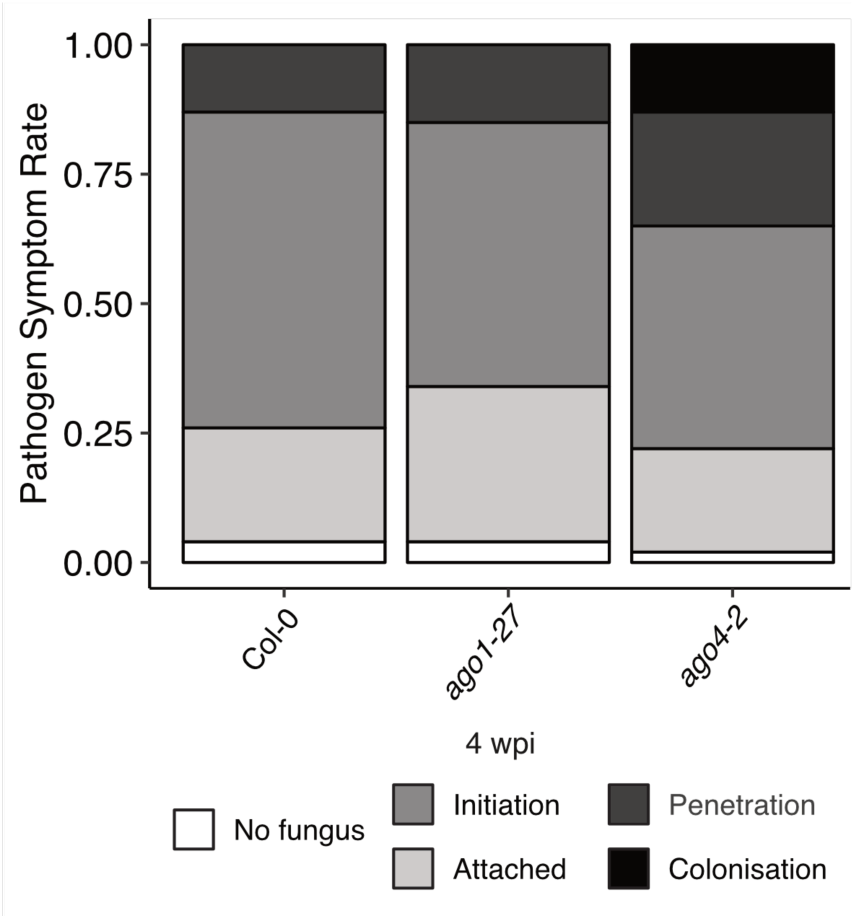
Colonization in *ago* mutants upon *T. thlaspeos* infection. Colonization was visualized after four weeks by staining with wheat germ agglutinin (WGA, fungal hyphae) and propidium iodide (PI, plant background). In at least 150 seedlings per line, fungal progression was classified into (i) attachment of the fungus to plant tissue, (ii) initiation of penetration as indicated by bulging of the hyphal tip, (iii) penetration into the plant tissue, and (iv) colonization along the vasculature. Similar results were obtained in three independent experimental replicates.

### AGO1 is a regulator of certain but not all bacterial infections

Motivated by the previous reports on the roles of AGO1 and AGO2 in immunity against bacterial pathogens (Ren et al., 2019; Zhang et al., 2011), we examined the roles of the selected *DCL* and *AGO* genes in infection by three different bacterial pathogens. Interestingly, AGO1 might be required for immunity against *X. fastidiosa*, since it was upregulated at late infection stages (Figure 4A), and both *ago1-27* and *ago1-46* displayed enhanced susceptibility (Figure 4B). All other tested genes showed wild type-like expression patterns and the respective mutants supported wild type-like infection success of *Pto* DC3000 (Supplemental Figure S3A and S3B) and *X. campestris pv. campestris* (Supplemental Figure S4A and S4B). The tested bacteria are Gram-negative γ-proteobacteria, including two belonging to the Xanthomonadaceae and colonizing xylem vessels (Supplemental Table S3). However, the positive regulatory function of AGO1 appears to be specific to immunity against *X. fastidiosa*.

**Figure 4:**
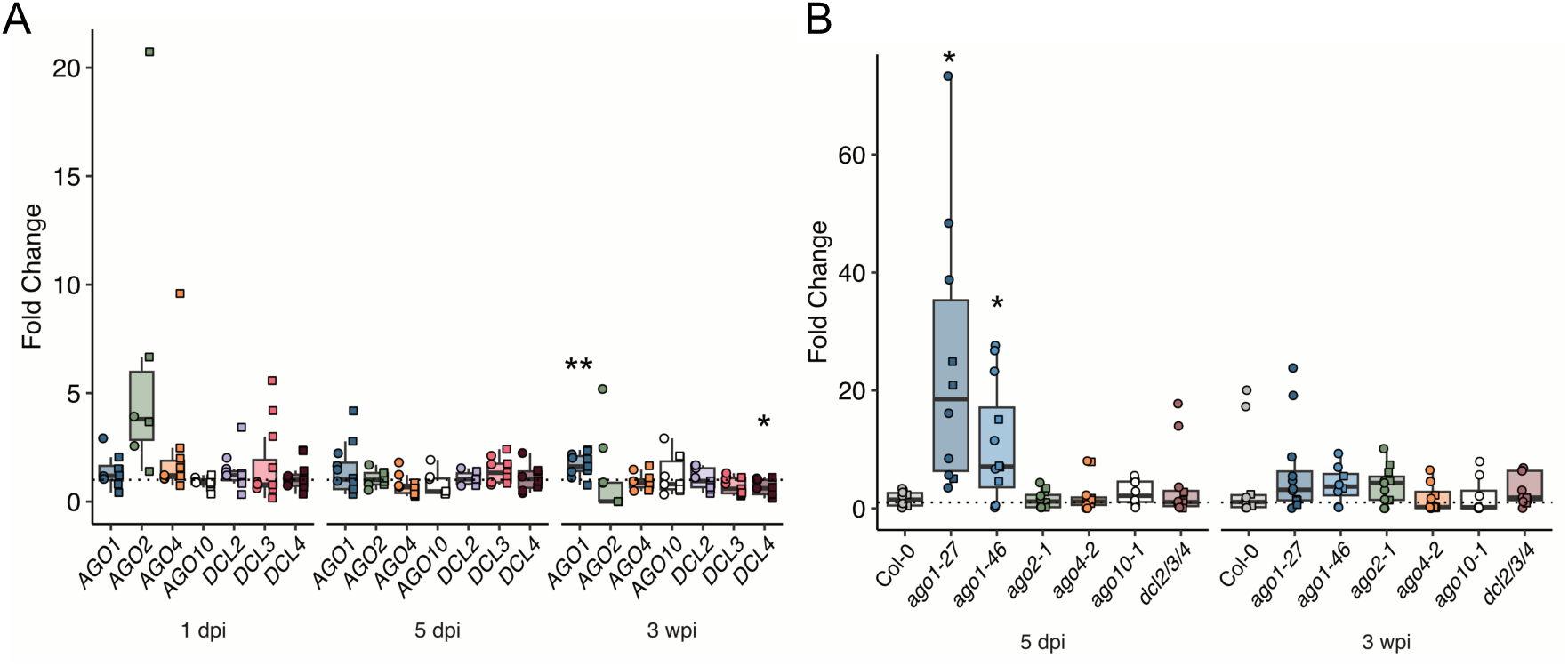
*DCL* and *AGO* gene expression patterns (A) and colonization in respective mutants (B) upon *X. fastidiosa subsp. fastidiosa* Temecula 1 infection. **(A)** Samples were collected from petioles at 1 (day post infection), 5 dpi, and 3 wpi (weeks post infection). The RNA levels are relative to mock and normalized against *CDKA*. The results of two independent biological replicates are depicted. **(B)** Pathogen load on *ago* and *dcl* mutants was assessed by RT-qPCR at 5 dpi and 3 wpi. The results of two independent biological replicates are depicted. Error bars show standard deviation. Statistical significance was assessed by two-sided Welch’s t-test (α = 0.05, p-values * < .05, ** < .01, *** < 0.001, **** < 0.0001). Symbols indicate number of independent biological replicate. Circle = first, square = second, diamond = third. The dashed line indicates a fold change = 1.

Previously, AGO2 and AGO4 were shown to positively regulate immunity against *Pto* DC3000 strains (Agorio & Vera, 2007; López et al., 2011; Zhang et al., 2011), which contrasts our findings. We neither detected any obvious induction of *AGO2* expression nor increased *Pto* DC3000 susceptibility in *ago2* and *ago4* mutants (Supplemental Figure S3). It is possible that the different outcomes for these mutants might depend on the methods of bacterial inoculation, as in the previous studies bacteria were applied by syringe-based leaf infiltration while in this study *Pto* DC3000 was sprayed onto the leaf surface. The two inoculation methods differ, as syringe inoculation bypasses stomatal immunity (Melotto et al., 2017).

### AGO2 and AGO10, but not AGO1, are potential regulators of fungal mutualism

Soybean AGO1 plays a positive role in the bacterial *Sinorhizobium* root-nodule symbiosis via ckRNAi (Ren et al., 2019). Furthermore, AGO1 has been speculated to regulate fungal symbiosis, supported by the prediction of plant mRNA targets of fungal sRNAs accumulating in the symbiosis between beneficial microorganisms and their hosts (Silvestri et al., 2019; Valdés-López et al., 2019; Wong-Bajracharya et al., 2022). Therefore, we next carried out colonization experiments with the mutualist basidiomycete *S. indica*, revealing *AGO4* downregulation at late time points (Figure 5A). The other tested *DCL* and *AGO* genes revealed no statistically significant changes in response to *S. indica* colonization at the investigated time points (Figure 5A). Interestingly, roots of *ago2-1* and *ago10-1* mutants showed a reduced colonization by *S. indica*, whereas *ago1-27*, *ago4-2* and *dcl2/3/4* exhibited wild type-like colonization (Figure 5B). It suggests that AGO2 and AGO10, but not AGO1, function as potential positive regulators during colonization in the mutualistic interaction of *A. thaliana* with *S. indica*, or negatively regulate immunity against this beneficial fungus. Moreover, although clade I AGO1 and AGO10 are phylogenetically related, AGO10 may have a specific function in fungal mutualism. Of note, miRNAs can be sequestered by different AGO proteins leading to different outcomes, e.g. as shown for miRNA165/166 in flower development, which depends on their binding to AGO1 and AGO10 (Ji et al., 2011).

**Figure 5:**
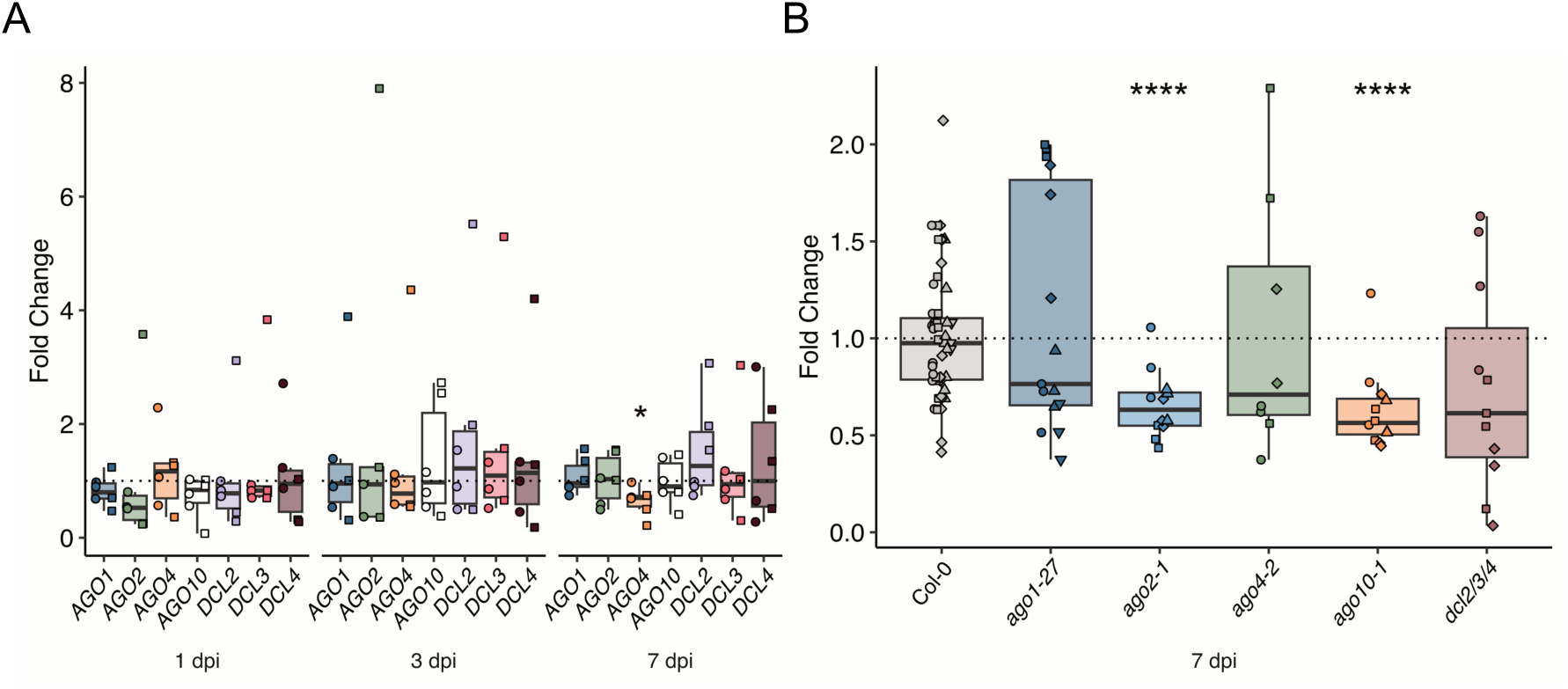
*DCL* and *AGO* expression patterns (A) and colonization in respective mutants (B) upon infection with *S. indica*. (**A)** Samples were collected from roots at 1 dpi (day post inoculation), 3 dpi, and 7 dpi. The RNA levels are relative to mock and normalized against *UBQ4*. The results of two independent biological replicates are depicted. **(B)** Pathogen load on *ago* and *dcl* mutants was assessed by measuring relative *S. indica* gDNA quantities with RT-qPCR at 7 dpi. The results of at least three independent biological replicates are depicted. Error bars show standard deviation. Statistical significance was assessed by two-sided Welch’s t-test (α = 0.05, p-values * < .05, ** < .01, *** < 0.001, **** < 0.0001). Symbols indicate number of independent biological replicate. Circle = first, square = second, diamond = third. The dashed line indicates a fold change = 1.

## Discussion

RNAi executed by DCL and AGO proteins is considered a conserved process regulating the outcome of plant-microbe interactions. Previous studies have described the roles of AGOs as both positive and negative regulators of plant immunity, likely linked to their binding of host endogenous or pathogen-derived sRNAs. Here, we i) confirm previous findings for AGO1 in negatively regulating immunity against *H. arabidopsidis* (Figure 1B) and ii) revealed a potential positive regulatory role of AGO1 in immunity against *X. fastidiosa* (Figure 4B). Moreover, we iii) identified clade I AGO1-related AGO10 and clade II AGO2 as positive modulators of *S. indica* root colonization (Figure 5B), and iv) revealed clade III AGO4 as a positive control element of *T. thlaspeos* infection (Figure 3). Thus, our broad scale phenotyping uncovered previously unknown positive and negative regulatory functions of different AGO proteins in the context of plant-microbe interactions.

AGO1’s negative adjustment of plant immunity is influenced by its role as a target for pathogen-derived sRNAs and its function in ckRNAi (Dunker et al., 2020; Weiberg et al., 2013). Therefore, it is possible that AGO1-related AGO10 and AGO2 might be hijacked by *S. indica*-secreted sRNAs, providing a possible molecular mechanism of their positive modulatory role in mutualism with this fungus. Indeed, the production of host and fungal-derived sRNAs has been revealed in the beneficial interaction of *Brachypodium distachyon* with *S. indica* (Šečić, Zanini, et al., 2021), which could result in PTGS of plant immunity genes, thereby facilitating *S. indica* colonization. This scenario is consistent with AGO1’s role in bacterial symbiosis, binding rhizobial tRNA-derived sRNA fragments (tRFs) that promote host nodulation (Ren et al., 2019). Additionally, clade I AGOs could influence the host’s transcriptional response to symbiosis, as reported for AGO5 in rhizobia-*Phaseolus vulgaris* symbiosis (Sánchez-Correa et al., 2022).

A positive immune regulatory function of AGO proteins has been linked to its binding of host endogenous sRNAs. For example, AGO2 binding of miR393b* triggers *MEMB12* (encoding a Golgi-localized SNARE protein) cleavage, which results in increased resistance due to the secretion and accumulation of the Pathogenesis-Related (PR) 1 protein (Zhang et al., 2011). It is possible that a potential function of AGO1 in restricting *X. fastidiosa* infection is related to endogenous host sRNAs associated with the control of PTI or its execution (Mitre et al., 2021; Navarro et al., 2006; Zhang et al., 2011). Given that ckRNAi functions in both directions (Cai et al., 2018), plants may also send AGO1-dependent sRNAs to alter *X. fastidiosa* growth, interfering with gene silencing in bacteria (Papenfort & Melamed, 2023).

The identification of AGO4 as a positive control element of resistance to *T. thlaspeos* expands the importance of this AGO protein beyond its requirement for immunity against *Pto* DC3000 and *B. cinerea* (Agorio & Vera, 2007; Rebolledo-Prudencio et al., 2022). Of note, upon infection with *Blumeria graminis* f.sp. *tritici*, *AGO4* was significantly downregulated in the wheat progenitor *Aegilops tauschii*, which was accompanied by a substantial reduction in AGO4a-sorted 24-nt siRNA levels, and enrichment for ‘response to stress’ gene functions, including receptor kinase, peroxidase, and pathogenesis-related genes, suggesting that AGO4 in some cases is a strong negative regulator of immunity (Geng et al., 2019). It further highlights the involvement of clade III AGOs-mediated TGS in the modulation of plant immunity. Interestingly, AGO1 was not required for *T. thlaspeos* infection at early stages of colonization. Therefore, other AGOs of clade I and clade II need to be investigated to address a putative role of PTGS in this fungal interaction during the established biotrophic phase or during fungal sporulation.

The other tested interactions did not reveal obvious phenotypes. This was unexpected given the functional conservation of AGOs, and the ubiquitous expression of key members like *AGO1* and *AGO4* in most tissues including leaves, roots, and the vasculature (Martín-Merchán et al., 2023; Wook et al., 2011). The transcriptional response of *DCL* and *AGO* genes was mostly not correlated with infection phenotypes in respective mutants. Considering the different infection types, from epidermal (*E. cruciferarum*), leaf mesophyll (*H. arabidopsidis*, *T. thlaspeos*, *Pto* DC3000), root (*S. indica*) to vascular tissues (*V. longisporum*, *X. campestris pv. campestris*, *X. fastidiosa*), a spatiotemporal resolution might be needed to observe changes in gene expression at the actual site of pathogen colonization (Dunker et al., 2020). We speculate that functional redundancy within the DCL and AGO family is likely accounting for wild type-like phenotypes in some of the tested interactions. This includes the *dcl2/3/4* triple mutant, therefore suggesting potential further redundancy of these three encoded proteins with DCL1. Moreover, infection with *E. cruciferarum* and *X. campestris pv. campestris* did not reveal infection phenotypes in the tested mutants (Figure 2B and Supplemental Figure S4B). This could suggest more functional redundancies among AGO proteins than expected.

To a large extent, the outcome of infection success is determined by the ability of the pathogen to suppress host immunity (Jones et al., 2024). This capacity is encoded in the pathogen’s repertoire of diverse molecular effectors (Y. Wang et al., 2022). For example, virulence of bacterial pathogens like *Pto* DC3000 and *X. campestris pv. campestris* mainly involves Type-3-secreted effector proteins (Y. Wang et al., 2022). *X. fastidiosa* lacks a Type-3 secretion system (Landa et al., 2022), and immune-suppressing protein effectors have not been described to date. Thus, the bacterium might not be able to overcome AGO1-mediated defences. Effector proteins have also been demonstrated to improve infection of powdery mildew fungi such as *E. cruciferarum* (Bourras et al., 2018). It is therefore possible that protein effectors could be largely responsible for virulence in a given microbe, contrasting to the virulence mechanisms of *B. cinerea* and *H. arabidopsidis*, which at least in part rely on sRNA-like effectors (Dunker et al., 2020; Weiberg et al., 2013). Moreover, protein effectors could suppress host RNAi (Hou et al., 2019; Navarro et al., 2006). Since pathotypes of microbial subspecies encode different effector repertoires, their interaction with the host’s RNAi machinery might differ (Qin et al., 2023; Weiberg et al., 2013).

To conclude, analysing the role of RNAi in plant immunity across taxonomically diverse microbes may be more complex and might need refined experimental set-ups beyond whole plant phenotyping with improved spatiotemporal resolution. We consider three levels of functional redundancy, which complicate the phenotypic analysis: i) similar or overlapping functions of DCL and AGO clade members, ii) microbial virulence conferred by protein effectors, including iii) microbial protein effectors suppressing host RNAi. Therefore, experimental studies would benefit from the use of higher order plant mutants (if not exhibiting severe developmental phenotypes), and combinatorial analysis with different pathotypes of microbial (sub-)species, as well as microbial mutants compromised e.g. for selected protein effectors or their secretion. Despite these considerations, our broad-scale phenotyping has uncovered previously unknown roles of AGO proteins.

## Material and methods

### Plant materials

*A. thaliana* Col-0 mutants included published *ago1-27, ago1-46, ago2-1, ago4-2, ago10-1,* and *dcl2/3/4* (Supplemental Table S1). Age-matched seeds were used for all experiments. The *eds1-2* mutant was used for propagation of the *E. cruciferarum* inoculum, and the *mlo2-5/6-2/12-1* mutant as an additional control for powdery mildew infection experiments (Consonni et al., 2006; Bartsch et al., 2006) .

### Primers

All primers used in this study are listed in Supplemental Table S3. To assess the expression levels of the selected *AGO/DCL* genes and also for quantification of microbial growth *in planta* (*H. arabidopsidis*, *X. fastidiosa*) by (RT)-qPCR, the *CDKA* gene expression was used as a reference. For *S. indica* colonization, *UBQ* was used as a reference gene.

### Microbial infections

#### Hyaloperonospora arabidopsidis

Plants for infection with *H. arabidopsidis* (GÄUM.) isolate Noco 2, *A. thaliana* plants were grown on soil under long day conditions (16 h light, 8 h dark, 60% relative humidity). Two weeks-old *A. thaliana* plants were inoculated with a final spore concentration of 2*10^4^ spores mL^-1^ as previously described (Ried et al., 2019). For biomass quantification, two leaves and two cotyledons were pooled for one replicate, followed by genomic DNA extraction with CTAB and RNase A treatment (Promega) (Chen and Ronald,1999). The isolated DNAs were diluted to 5 ng/µl. *H. arabidopsidis* gDNA relative to *A. thaliana* was quantified by qPCR with Primaquant low ROX qPCR master mix (Steinbrenner Laborsysteme) according to the manufacturer’s instructions (95 °C 3 min, 95 °C 20 s, 60 °C 30 s, 72 °C 40 s, 40 cycles, and subsequent melting curve analysis.

For RT-qPCR, four leaves were pooled for one replicate. The CTAB method was used for total RNA extraction (Bemm et al., 2016). Genomic DNA was removed by DNase I digestion (Sigma-Aldrich) following the manufacturer’s instructions. For cDNA synthesis with the Maxima H Minus Reverse Transcriptase (Thermo Fisher Scientific) kit, 1 μg of total RNA from each sample was used. Relative gene expression was quantified by qPCR with the Primaquant low ROX qPCR master mix (Steinbrenner Laborsysteme), according to the manufacturer’s instructions (95 °C 3 min, 95 °C 20 s, 60 °C 30 s, 72 °C 40 s, 40 cycles, melting curve analysis).

#### Erysiphe cruciferarum

Plants for *E. cruciferarum* infection were grown on SoMi 513 soil (Hawita, Vechta, Germany) in 9*9 cm pots under short-day conditions with an 8-h photoperiod at 22 °C and 16 h darkness at 20° C. *E. cruciferarum* (in-house isolate of RWTH Aachen) was cultivated selectively on *A. thaliana eds1*-2 (Bartsch et al., 2006) at 20 °C with an 8-h photoperiod. Spores from three pots of plants, collected at 20-28 dpi, were used for inoculation of 10 pots. For this, four weeks-old healthy Col-0 plants, the selected mutant lines and the resistant *mlo2-5/6-2/12-1* triple mutant (negative control) were placed in an inoculation tower and heavily infected inoculum plants were gently agitated to release spores. To determine fungal entry rates, leaves were harvested at 48 hours post inoculation (hpi) and collected in 80% EtOH for de-staining of leaf pigments. Fungal structures were stained with Coomassie staining solution (45% MeOH (v/v), 10% acetic acid (v/v), 0.05% Coomassie blue R-250 (w/v)). Samples were double-blinded and leaves were analysed by light microscopy. The fungal entry rate was determined as the percentage of spores successfully developing secondary hyphae over all spores that attempted penetration, visible by the presence of an appressorium (Kusch et al. 2019). At least 100 interaction sites on leaves of three different plants per independent replicate were analysed.

For RT-qPCR, total RNA was extracted from leaves of uninoculated Arabidopsis and leaves sampled at 8, 48, or 96 hpi with *E. cruciferarum* using TRI reagent ® (Sigma Aldrich), each from one leaf of three different plants per sample. The remaining DNA was digested using DNase I (Thermo Fisher Scientific, USA). For cDNA synthesis 1 µg RNA was used with the High-Capacity cDNA Reverse Transcription Kit (Applied Biosystems). Relative expression of target genes was quantified by RT-qPCR (95 °C 3 min, 95 °C 10 s, 60 °C 60 s; 40 cycles; melting curve analysis) with the Takyon no ROX SYBR 2X master mix (Eurogentec).

#### Verticilium longisporum

Arabidopsis mutants used for *V. longisporum (VL43)* (Zeise & Von Tiedemann, 2002)infection were grown directly on soil in a climate chamber with 22 °C/18 °C day/night cycle with 8 h of light. For infection of mutant lines, an inoculation suspension was used. This suspension was prepared by flooding a fully grown three weeks-old culture of *V. longisporum* grown on Potato Dextrose Agar (PDA) agar (Carl Roth, Art. No. X931.1) in a petri dish at 22 °C, in the dark with 10 mL ddH_2_O. The petri dish was scraped with a small metal spatula to release conidia in suspension. The suspension was filtered through miracloth (Calbiochem, 475855) to exclude mycelium and spore concentration was determined by using a hemocytometer (Thoma counting chamber - Marienfeld). The final concentration of the spore suspension was adjusted to 10.000.000 spores/mL. Two weeks-old seedlings of *A. thaliana* (Col-0 and mutant lines) were infected by pipetting 1 mL of inoculation suspension directly in the soil. Plant material was harvested 1, 7 and 35 dpi and ground on liquid nitrogen. Total RNA extraction was done by using TRIzol® (Invitrogen) and Zymo RNA Clean & Concentrator-25 Kit with in-column DNase I digestion. cDNA synthesis was generated using 200 ng/µL of the extracted total RNA using RevertAid Reverse transcriptase (Thermo Scientific) following manufacturer guidelines. For subsequent qPCR the SYBR™ Green PCR Master Mix (Applied Biosystems) was used (Thermo Fisher Scientific 4309155). For this, 10 µl reactions were set up, consisting of 5 µl SYBR master mix, 0.5 µL of each primer, 3.5 µL H_2_0 and 0.5 µL cDNA, run with 3 min of 95 °C followed by 40 cycles of 95 °C for 10s, 60 °C for 1 min, and subsequent melting curve analysis, on a QuantStudio 6 Flex (Applied Biosystems).

#### Thecaphora thlaspeos

One sterilized seed of an Arabidopsis line was co-germinated with 300 sterilized teliospores of *T. thlaspeos* (collection 2022, Frantzeskakis et al., 2017) in 300 µL liquid half-strength Murashige and Skoog with Nitrate (MSN) medium (Duchefa) containing 1% sucrose in a well of a 96-well plate. The infections were incubated for four weeks in a light chamber for *A. thaliana* at long-day conditions (120 μE, 12 h 21 °C light, 12 h 18 °C darkness). Seedlings were then stained with wheat germ agglutinin (WGA) and propidium iodide (PI) as done previously described (Frantzeskakis et al., 2017) and scored microscopically for fungal infection stages (Zeiss Axio Immager M1). Up to 160 seedlings were inspected per line and experiment.

#### Serendipita indica

*A. thaliana* mutant lines were grown on vertical square Petri dishes on *Arabidopsis thaliana* Salt medium (ATS) (Lincoln et al., 1990) without sucrose and supplemented with 4.5 g/L Gelrite (Duchefa #G1101) in a 22 °C day/18 °C night cycle (8 h of light). Spores of *S. indica* (IPAZ-11827, Institute of Phytopathology, Giessen, Germany) were freshly isolated from the plates by scrapping the agar using water supplied with 0.002% Tween 20 added and then filtered through miracloth (Merck Millipore), centrifuged at 3.000 x g for 7 min, then resuspended in water supplied with 0.002% Tween 20 and adjusted to 500.000 spores mL^-1^.

Roots of 14 days-old plants were inoculated with 1 mL of a suspension of 500.000 chlamydospores mL^-1^ in water with 0.002% Tween 20 per Petri dish. Control plants were treated with water supplied with 0.002% Tween 20 (mock). Inoculated roots of different mutants are harvested after seven days, and ground for 1 min at 30 Hz with the pre-cooled Retsch Mill (Tissue Lyser II, Retsch, Qiagen). For quantification of *S. indica* colonization, genomic DNA was extracted using a Qiagen DNA extraction kit (QIAGEN, 69504). Fungal colonization was quantified using internal transcribed spacer (ITS) primers (see Supplemental Table 3) and SYBR Green JumpStart Taq ReadyMix (Sigma Aldrich, 1003444642) with a QuantStudio5 Real-Time PCR System (Applied Biosystems). 2 µL ROX (CRX reference dye, Promega, C5411) were added to 1 mL SybrGreen as a passive reference dye that allows fluorescent normalization for qPCR data. The PCR conditions were 95 °C for 5 min followed by 40 cycles of 95 °C for 15 s, 60 °C for 30 s, and 72 °C for 30 s followed by melting curve analysis.

For *DCL* and *AGO* gene expression, Arabidopsis Col-0 plants were grown on ATS plates and inoculated with *S. indica* spores as previously described above. Control plants were treated with water containing 0.002% Tween 20 (mock). Inoculated roots were harvested at 1, 3, and 7 dpi, ground with the tissue lyser, and RNA was extracted using Trizol and Zymo kit (Zymo research R2070), with a subsequent in-column DNase digestion. cDNA was generated from 1 µg RNA using Revert Aid Reverse transcriptase. Gene transcription was quantified by qPCR using SYBR Green JumpStart Taq ReadyMix (Sigma Aldrich, 1003444642) with QuantStudio5 Real-Time PCR System (Applied Biosystems). 2 µL ROX (CRX reference dye, Promega, C5411) were added to 1 mL SybrGreen as a passive reference dye that allows fluorescent normalization for qPCR data. The PCR conditions were 95 °C for 5 min followed by 40 cycles of 95 °C for 15 s, 60 °C for 30 s, and 72 °C for 30 s followed by a melting curve analysis. *Ubiquitin* (*UBQ4, AT5G20620*) was used as a housekeeping gene for all experiments. Roots from two ATS plates were harvested and considered as one biological replicate. The results of three or more independent biological replicates are included in the data analysis.

#### *Pseudomonas syringae* pv*. tomato* DC3000

Plants for *Pto* DC3000 infection were grown on soil with 10 h light and 55% humidity for four to five weeks. *Pto* DC3000 was routinely grown at 28 °C on King’s B plates with 1% Agar. Overnight plate-grown *Pto* DC3000 cells were resuspended in 10 mM MgCl_2_ and 0.04% Silwet L-77 and diluted to OD_600_ = 0.02. The *A. thaliana* plants were sprayed from below and on top with inoculum. Discs of the infected leaves (one disc per leaf, 0.6 cm diameter) were excised at 1 dpi and 3 dpi. Four leaf discs from one plant were pooled and ground in 200 µL 10mM MgCl_2_. Serial dilutions were plated on King’s B medium with rifampicin (50 μg mL^-1^) and bacterial colonies were quantified after two days of incubation at 28 °C. At least four plants per genotype and time points were harvested and plated. Results of three independent rounds of infection are included.

For qRT-PCR, two inoculated or mock-treated (sprayed with buffer-only) leaves were harvested at 6 hpi, 1 dpi and 3 dpi. Leaf material was ground using a tissue lyser and RNA extractions performed with Trizol reagent (Invitrogen, USA) according to the manufacturer’s protocol and the Zymo RNA Clean & Concentrator Kit, including in-column DNase treatment. RT-qPCR was performed using the NEB Luna^®^ Universal One-Step RT-qPCR Kit (E3005) according to the manufacturer’s instructions (55 °C 10 min, 95 °C 1 min, 95 °C 10 s, 60°C 30 s, 45 cycles, and subsequent melting curve analysis). Reactions were set-up in duplicates using 10 ng RNA in 10 µl reactions. At least four samples per time point and treatment were analysed, two rounds of infection are included.

#### *Xanthomonas campestris* pv*. campestris*

Plants for *X. campestris* pv*. campestris* infection were grown on soil with 10 h light and 55% humidity for four to five weeks. *X. campestris* pv*. campestris* 8004 was routinely grown at 28 °C on NYG (Nutrient Yeast Glycerol Agar, Daniels et al., 1984) media with 1% agar. Inoculum was prepared freshly by scraping bacteria from plates and resuspended in 1x PBS for a final OD_600_ of 0.4. Four leaves per plant were inoculated by application of 5 µl drops of bacterial suspension onto the midvein of leaves prior to pricking with a 0.4 * 20 mm needle five times. Plants were covered in a plastic bag for the first two days to create optimal infection conditions with high humidity. Discs of the inoculated leaves (one disc per leaf, 0.6 cm diameter) were excised at 3 dpi and 5 dpi. Two leaf discs were pooled and ground in 200 µL 10 mM MgCl_2_. Serial dilutions were plated on King’s B medium supplemented with rifampicin (50 μg mL^-1^), and bacterial colonies were quantified at two days after incubation at 28 °C. Suspensions that resulted in no colonies were excluded from the analysis. At least four samples per genotype and time point were harvested and plated. Results of three independent rounds of infection are included.

For qRT-PCR, two inoculated or mock-treated leaves were harvested at 1 dpi, 3 dpi and 7 dpi. RNA extraction and qRT-PCR reactions were performed as described above for *Pto* DC3000.

#### *Xylella fastidiosa* subsp. *fastidiosa*

Plants for *X. fastidiosa* subsp. *fastidiosa* infection were grown on soil with 10 h light and 55% humidity for four to five weeks. *X. fastidiosa* subsp. *fastidiosa* Temecula 1 (ATCC 700964) was routinely grown at 28 °C on PD3 plates (Pierce’s Disease 3, Davis et al. 1981) for approx. seven to ten days. The inoculum was prepared freshly by scraping bacteria from plate and resuspending it in 1x PBS for a final OD_600_ of 0.5. Four leaves per plant were inoculated by application of 5 µl drops of bacterial suspension onto the midvein of leaves prior to pricking with a 0.4 * 20 mm needle 5 times. Two petioles were combined and harvested at 5 dpi and 3 weeks post inoculation (wpi). RNA was extracted from 2 petioles of infected samples after disruption with a tissue lyser, using Trizol reagent (Invitrogen, USA) according to the manufacturer’s protocol and Zymo RNA Clean & Concentrator Kit, including in-column DNase treatment. qRT-PCR was performed with 10 ng RNA using the NEB Luna^®^ Universal One-Step RT-qPCR Kit (E3005), in 10 µL reactions according to manufacturer guidelines (55 °C 10 min, 95 °C 1 min, 95 °C 10 s, 60 °C 30 s, 45 cycles, and subsequent melting curve analysis) using primers for *Xf*16S and *CDKA* (see Supplemental Table S3) to normalize for plant material. At least four samples per genotype and time points were analysed. Results of two independent rounds of infections are included. For qRT-PCR, two inoculated leaves or mock-treated leaves were harvested at 1 dpi, 5 dpi and 3 wpi. RNA extraction and qRT-PCR reactions were performed as described above for *Pto* DC3000.

### Statistical analysis

All data was analysed using R (version 2023.06.0+421) and statistical analysis was performed using the stats-package (*R: The R Project for Statistical Computing*, n.d.). For infection data, mutant measurements were compared to respective measurements in Col-0. For RT-qPCR and qPCR data analysis, expression values were analysed using the 2^(-ΔΔct)^ method (Livak & Schmittgen, 2001) and normalized against *CDKA* or *UBQ* (*S. indica*) as housekeeping genes and the average of respective mock samples. For *E. cruciferarum,* all time points were compared to T0. Significance was assessed by two-sided Welch’s t-test (α = 0.05, p-values * < .05, ** < .01, *** < 0.001, **** < 0.0001) using the stats-package in R.

### Use of public data

The heatmap showing the differential expression of *DCL* and *AGO* genes (Supplemental Figure S1) was generated using data from Bjornson et al. Nat Plants 2021 (Bjornson et al., 2021) (Supplemental Table S1 and S2) with Python v3.11.4 (Guido van Rossum & Fred L. Drake, Jr., n.d.) and Seaborn v0.12.2 (Waskom, 2021).

### Accession numbers

Genes reported in this article can be found in the GenBank/RGAP databases under the following accession numbers: *AGO1 (AT1G48410), AGO2 (AT1G31280), AGO4 (AT2G27040), AGO10 (AT5G43810), DCL2 (AT3G03300), DCL3 (AT3G43920), and DCL4 (AT5G20320), CDKA (AT3G48750.1), UBQ4 (AT5G20620)*.

## Supplemental data files

Supporting information Supplemental Figures S1-4 and Supplemental Tables S1-3 are available online.

**Supplemental Figure S1:** Differential expression of *DCL* and *AGO* genes after treatment with different MAMPs using publicly available results of a large scale RNAseq experiment (Bjornson et al., 2021).

**Supplemental Figure S2:** *DCL* and *AGO* gene expression patterns upon *V. longisporum* infection.

**Supplemental Figure S3:** *DCL* and *AGO* gene expression patterns (A) and colonization in respective mutants (B) upon *Pseudomonas syringae* pv*. tomato* (*Pto*) DC3000 infection.

**Supplemental Figure S4:** Expression of *A. thaliana DCL* and *AGO* genes (A) and colonization in respective mutants (B) upon *Xanthomonas campestris* pv*. campestris* 8004 infection.

**Supplemental Table S1:** Overview of investigated genes, their corresponding mutants used in this study, and their roles in RNAi and plant immunity

**Supplemental Table S2:** Overview of microorganisms used in this study

**Supplemental Table S3:** Overview of primers used in this study

## Author contributions

A.R., H.T., S.N., B.L., S.F., J.K., J.S., E.S., S.K., V.G., M.F., A.W., K.H.K., R.P., S.R. designed the methodology; A.R., H.T., S.N., B.L., S.F., T.A., J.P., E.S. performed research; A.R., H.T., S.N., B.L., S.F., V.G., P.B., K.B. analysed data; A.G., J.K., V.G., A.W., K.H.K., R.P., S.R. conceived the project and supervised the research; A.R., H.T., S.N., S.R. wrote the manuscript with input from all authors.

## Acknowledgements

This work was enabled by grants within the research unit FOR5116 “exRNA” funded by the Deutsche Forschungsgemeinschaft (DFG). Individual grants of this research unit comprise DFG projects FE448/15-1 to M.F, GO 2037/8-1 to A.G, GO 2064/4-1 to V. G., KE 856/8-1 to J.K., KO 1208/32-1 to K.K., PA 861/22-1 to R.P, RO 3550/16-1 to S.R. and WE 5707/2–1 to A.W. S.N. was supported by a Dr. Ernst-Leopold Klipstein Foundation grant. We thank Elif Olkun for technical support.

