## Supplemental data for "Broad-scale phenotyping in Arabidopsis reveals varied involvement of RNA interference across diverse plant-microbe interactions"

**Supplemental materials**

**Supplemental Figure S1:** Differential expression of *DCL* and *AGO* genes after treatment with different MAMPs using publicly available results of a large scale RNAseq experiment (Bjornson et al. 2021).

**Supplemental Figure S2:** *DCL* and *AGO* gene expression patterns upon *V. longisporum* infection.

**Supplemental Figure S3:** *DCL* and *AGO* gene expression patterns (A) and colonization in respective mutants (B) upon *Pseudomonas syringae* pv. *tomato* (*Pto*) DC3000 infection.

**Supplemental Figure S4:** Expression of *A. thaliana DCL* and *AGO* genes (A) and colonization in respective mutants (B) upon *Xanthomonas campestris* pv. *campestris* 8004 infection.

**Supplemental Table S1:** Overview of investigated genes, their corresponding mutants used in this study, and their roles in RNAi and plant immunity

**Supplemental Table S2:** Overview of microorganisms used in this study

**Supplemental Table S3:** Primers used in this study

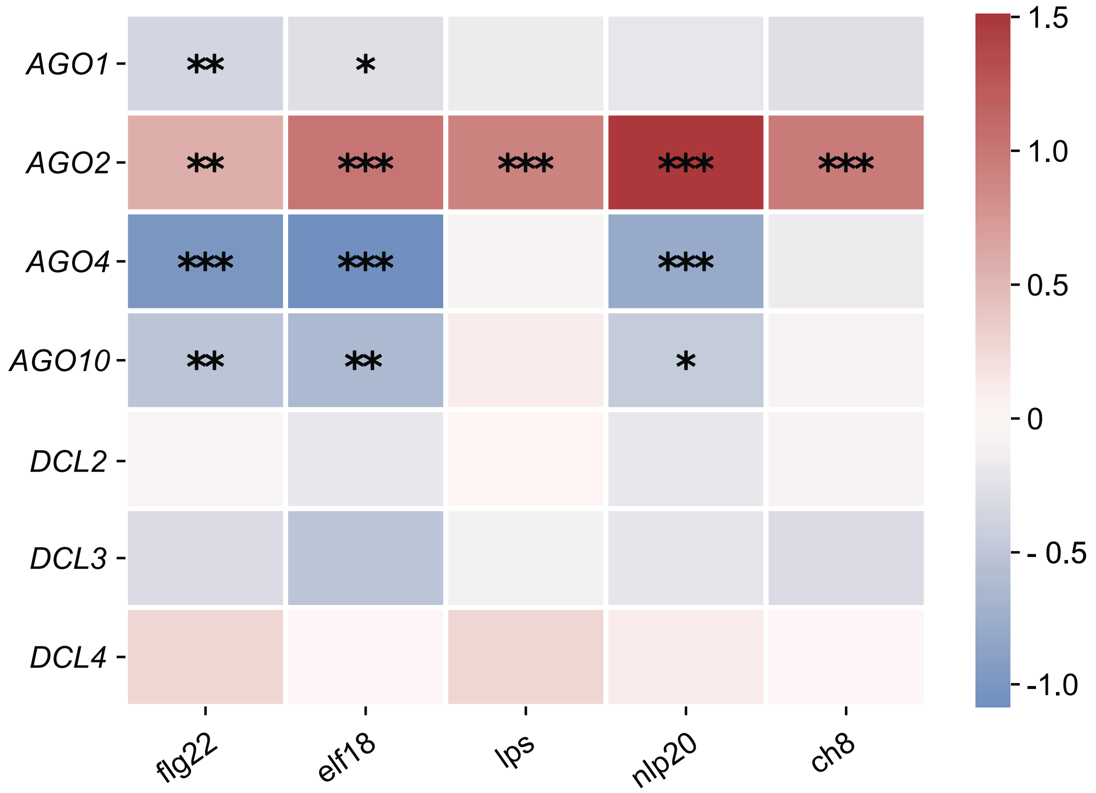


**Supplemental Figure S1: Differential expression of *DCL* and *AGO* genes after treatment with different MAMPs using publicly available results of a large scale RNAseq experiment** (Bjornson et al. 2021)**.** Data were retrieved from a publicly available large-scale RNAseq experiment (Bjornson et al. Nat Plants 2021) and are based on the treatment of Col-0 with the bacterial MAMPs flg22 (peptide epitope of bacterial flagellin), elf18 (peptide epitope derived from bacterial elongation factor Tu) and LPS-associated 3OH-FA (bacterial hydroxylated fatty acid, lipo-polysaccharide), the universal nlp20 peptide (bacterial, oomycete and fungal peptide) as well as the fungal MAMP CO8 (chitooctaose, fragment of the fungal cell wall). The published DESeq2 reports were mined for MAMP-responsive expression of genes of interest (*AGO1, AGO2, AGO4, AGO10, DCL2, DCL3* and *DCL4*). Depicted is the fold change of the indicated genes (log2) at 90 min after MAMP treatment. Red colour represents upregulation and blue colour downregulation of respective genes and significant differences are indicated with asterisks ($\alpha = 0.05$, adjusted p-values * < .05, ** < .01, *** < 0.001).

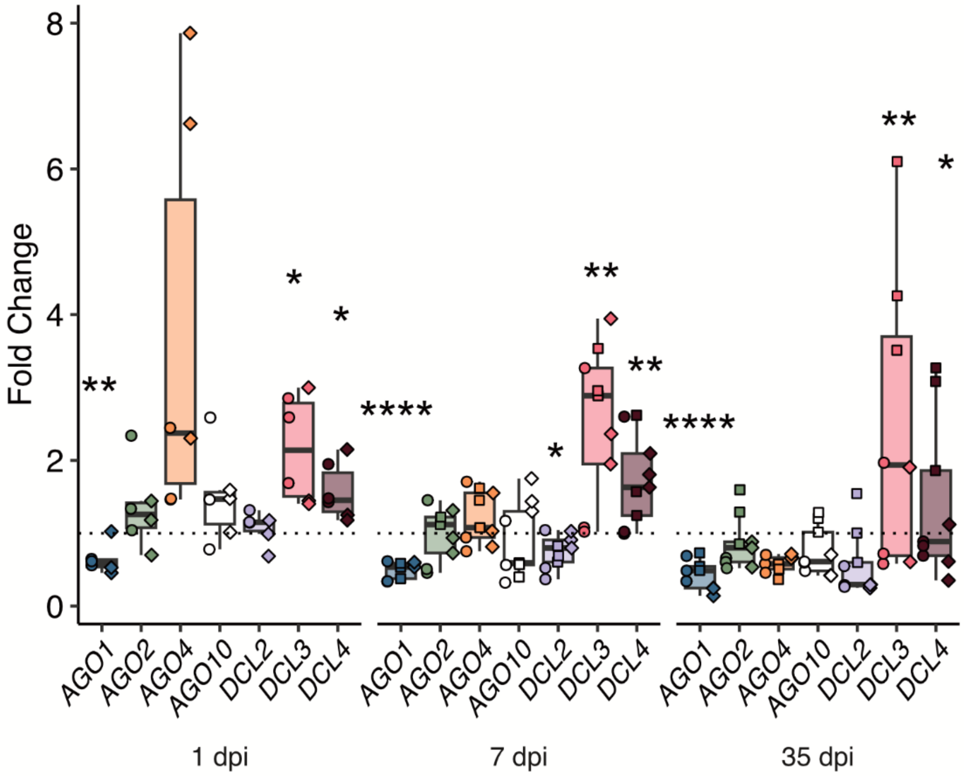


**Supplemental Figure S2: *DCL* and *AGO* gene expression patterns upon *V. longisporum* infection.** Samples were collected at 1 (days post inoculation), 7 dpi, and 35 dpi. The RNA levels are relative to mock treatment and normalized against *CDKA*. The results of three independent biological replicates are depicted. Error bars show standard deviation. Statistical significance was assessed by two-sided Welch's t-test (α = 0.05, p-values * < .05, ** < .01, *** < 0.001, **** < 0.0001). Symbols indicate the number of independent biological replicates. Circle = first, square = second, diamond = third. The dashed line indicates a fold change = 1.

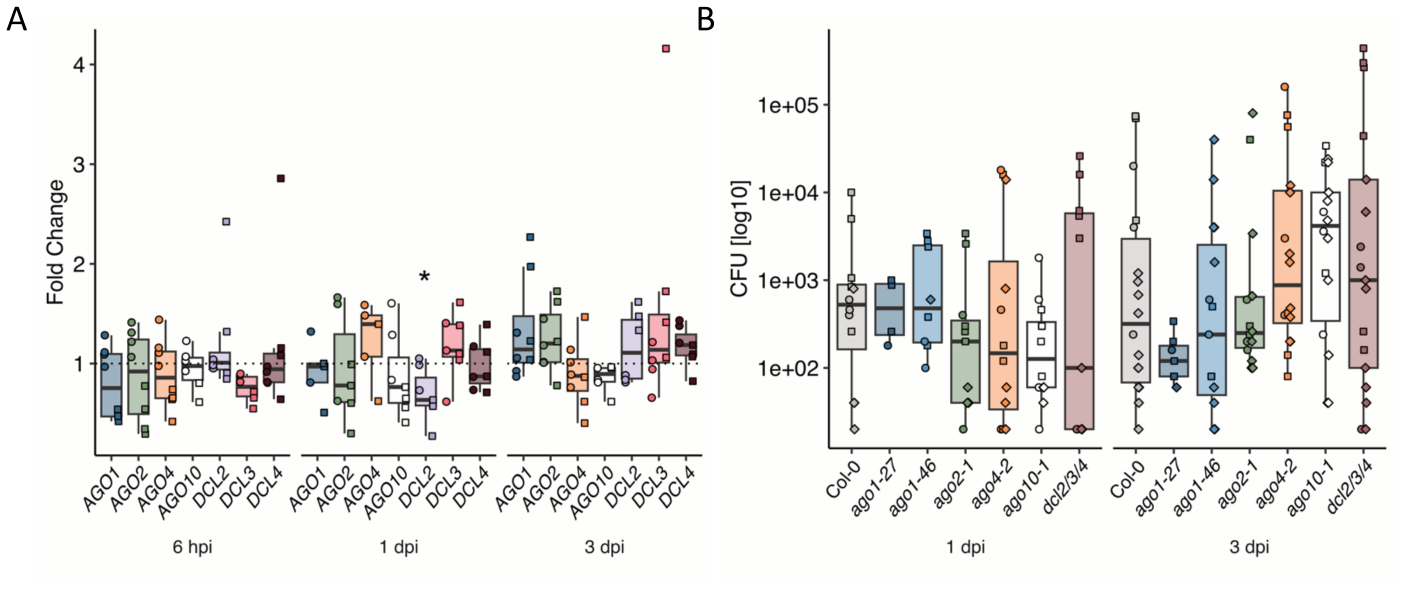


**Supplemental Figure S3:** ***DCL*** **and** ***AGO*** **gene expression patterns (A) and colonization in respective mutants (B) upon** ***Pseudomonas syringae*** **pv.** ***tomato*** **(*****Pto*****) DC3000 infection.** **(A)** Samples were collected from leaf material at 6 hpi (hours post infection), 1 dpi (day post infection), and 3 dpi. The RNA levels are relative to mock and normalized against *CDKA*. The results of two independent biological replicates are depicted. **(B)** Pathogen load on *ago* and *dcl* mutants was assessed by determining the number of colony-forming units at 1 dpi and 3 dpi. The results of three independent biological replicates are depicted. Error bars show standard deviation. Statistical significance was assessed by two-sided Welch's t-test (α = 0.05, p-values * < .05, ** < .01, *** < 0.001, **** < 0.0001). Symbols indicate number of independent biological replicate. Circle = first, square = second, diamond = third. The dashed line indicates a fold change = 1.

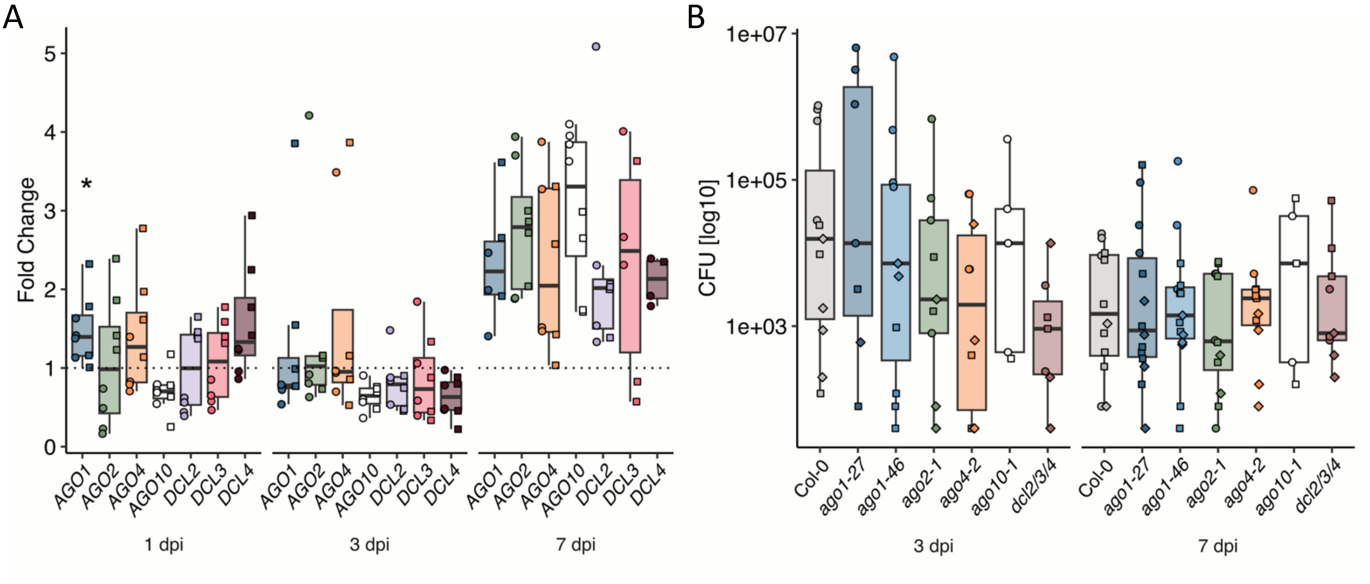


**Supplemental Figure S4: *DCL* and *AGO* gene expression patterns (A) and colonization in respective mutants (B) upon *Xanthomonas campestris* pv. *campestris* 8004 infection. (A)** Samples were collected from leaf material at 1 (day post infection), 3 dpi, and 7 dpi. The RNA levels are relative to mock and normalized against *CDKA*. The results of two independent biological replicates are depicted. **(B)** Pathogen load on *ago* and *dcl* mutants was assessed by determining the number of colony-forming units at 3 and 7 dpi. The results of three independent biological replicates are depicted. Error bars show standard deviation. Statistical significance was assessed by two-sided Welch's t-test ($\alpha = 0.05$, p-values * < .05, ** < .01, *** < 0.001, **** < 0.0001). Symbols indicate number of independent biological replicate. Circle = first, square = second, diamond = third. The dashed line indicates a fold change = 1.

**Supplemental Table S1:** Overview of investigated genes, their corresponding mutants used in this study, and their roles in RNAi and plant immunity.

| Protein/gene | Mutant | Role in RNAi | Role in immunity | References |
|---|---|---|---|---|
| AGO1 (*AT1G48410*) | *ago1-27*<br>*ago1-46* | • mi/siRNA-directed gene silencing<br>• PTGS | • Plays a role in PTI against bacterial pathogens; regulates response to viral infections | Qu, Ye, and Morris 2008; Morel et al. 2002; Zhang et al. 2018 |
| AGO2 (*AT1G31280*) | *ago2-1* | • Playing a significant role in antiviral RNAi | • Modulates defence against RNA viruses and regulates immune responses against viruses<br>• Anti-fungal immunity | Carbonell 2017; Harvey et al. 2011; Jaubert et al. 2011 |
| AGO4 (*AT2G27040*) | *ago4-2* | • Mediates TGS through siRNA-guided DNA methylation | • Plant defence against pathogens<br>• Anti-bacterial immunity<br>• Anti-viral immunity | Zhang et al. 2018; Agorio and Vera 2007; Brosseau et al. 2016 |
| AGO10 (*AT5G43810*) | *ago10-1* | • Regulates shoot apical meristem development<br>• PTGS | • Implicated in defence against fungal pathogens and symbiosis<br>• Anti-viral immunity | Harvey et al. 2011; Carbonell et al. 2012; Mallory et al. 2009 |
| DCL2 (*AT3G03300*) | *dcl2/3/4* | • Processes dsRNAs into 22-nt siRNAs | • Regulates defence responses against RNA viruses | Bouché et al. 2006; Zhang et al. 2018 |
| DCL3 (*AT3G43920*) | *dcl2/3/4* | • Generates 24-nt siRNAs, playing a major role in RdDM and TGS | • Essential for defence against DNA viruses and chromatin modifications | Singh et al. 2019; Vaucheret 2006; Xie et al. 2004 |
| DCL4 (*AT5G20320*) | *dcl2/3/4* | • Produces 21-nt siRNAs, crucial for PTGS | • Plays a role in defence against RNA viruses and response to abiotic stresses | Qu, Ye, and Morris 2008; Deleris et al. 2006; Gasciolli et al. 2005 |

**Supplemental Table S2:** Overview of microorganisms used in this study.

| Microbe | Lifestyle | Molecular patterns | Host target tissue | Infection / colonization strategy | References |
|---|---|---|---|---|---|
| *Erysiphe cruciferarum* | • Obligate biotroph | • chitin | • Leaf epidermal cell layer | • Forming appressoria<br>• Formation of haustoria | Glawe 2008; O'Connell and Panstruga 2006; Spanu et al. 2010 |
| *Hyaloperonospora arabidopsidis* Noco 2 | • Obligate biotroph | • β-glucans<br>• eicosapolyenoic acids<br>• nlp20 | • Lower leaf surface;<br>• Leaf epidermal cells | • Penetrates through the cuticle<br>• Formation of intercellular hyphae and haustoria | Dunker et al., 2019 |
| *Pseudomonas syringae* pv. *tomato* DC3000 | • Epiphyte and endophyte | • 3OH-FA<br>• csp22<br>• elf18 (EF-Tu);<br>• flg22<br>• PGN | • Leaf surface<br>• Leaf apoplast | • Entering into plant cells via stomata/ wounds<br>• Growth in apoplast leading to leaf spots | Xin and He 2013 |
| *Serendipita indica* | • Endophyte Biotrophic Saprophyt e | • chitin | • Root epidermal tissue<br>• Root cortical tissue<br>• Root hairs | • Fungal hyphal formation and epidermal cell penetration<br>• Colonization of the intercellular space | Deshmukh et al. 2006; Qianget al. 2011; Osborne et al. 2023 |
| *Thecaphora thlaspeos* | • Biotroph | chitin | • Stems<br>• Along vascular tissue | • Penetrating through natural openings like stomata or wounds | Courville et al. 2019; Frantzeskak is et al. 2017; Plücker et al. 2021 |

| | | | | | |
|---|---|---|---|---|---|
| *Verticillium lonngisporum* (*Vl43*) | • Hemi-biotroph | • β-glucans<br>• chitin | • Root and stem tissues<br>• Vasculature (xylem) | • Infects plants through the roots<br>• Penetrates root tissues and spreads systemically through the xylem vessels<br>• Produces microsclerotia | Liu et al. 2014 |
| *Xanthomonas campestris* pv. *campestris* 8004 | • Epiphyte and endophyte<br>• Biotroph and necrotroph | • 3OH-FA<br>• csp22<br>• elf18<br>• flg22<br>• PGN | • Vasculature (xylem)<br>• Apoplast | • Infection from seed material<br>• Entering adult plants via wounding, hydathodes, and roots<br>• Growth in vascular tissue<br>• Chlorotic/ necrotic lesions extending from leaf margins | Luneau et al. 2022 |
| *Xylella fastidiosa* subsp. *fastidiosa* Temecula 1 | • Endophyte | • elf18/26<br>• csp22<br>• EPS<br>• LPS<br>• PGN | • Vasculature (xylem) | • Transmission via insect vector results in drought-like symptoms | Mitre et al. 2021; Roper, Castro, and Ingel 2019; Landa et al. 2022 |

**Supplemental Table S3:** Overview of primers used in this study

| Primer | Sequence |
|---|---|
| Xf16S | FW: ACCTGGTCTTGACATCTGCG<br>RV: CACCATTACGTGCTGGCAAC |
| HaACT | FW: GTGTCGCACACTGTACCCATTTAT<br>RV: ATCTTCATCATGTAGTCGGTCAAGT |
| ITS | FW: CAACACATGTGCACGTCGAT<br>RV: CCAATGTGCATTCAGAACGA |
| UBI4 | FW: GCTTGGAGTCCTGCTTGGACG<br>RV: CGCAGTTAAGAGGACTGTCCGGC |
| CDKA | FW: ATTGCGTATTGCCACTCTCATAGG<br>RV: TCCTGACAGGGATACCGAATGC |
| AGO2 | FW: TGCAGAAGCTCATCTTCGAG<br>RV: GCGGCTGCTTAAAGTTCTTC |
| AGO4 | FW: GCCGATCTGCTATGCTCACT<br>RV: GGCGACGTTGTCTTTGAGTC |
| AGO10 | FW: GTCTCTATAGTTCCTCCAGCG<br>RV: TCCTTCAAGGCTGGTAAAGG |
| DCL2 | FW: GGTTGGATGGTTCCAGGTCA<br>RV: ACAACATCGGCCACACTCTT |
| DCL3 | FW: AGCCGTTGCTTTCTCCACTT<br>RV: GTGGCTTGTGCTTTGACACC |
| DCL4 | FW: AGGCTGCACAGCTGATGATT<br>RV: TCCAGCCCAACAAGATCCAC |

Dunker, Florian, Adriana Trutzenberg, Jan S Rothenpieler, Sarah Kuhn, Reinhard Pröls, Tom Schreiber, Alain Tissier, et al. n.d. 'Oomycete Small RNAs Bind to the Plant RNA-Induced Silencing Complex for Virulence'. *eLife* 9:e56096. https://doi.org/10.7554/eLife.56096.

Frantzeskakis, Lamprinos, Kaitlyn J. Courville, Lesley Plücker, Ronny Kellner, Julia Kruse, Andreas Brachmann, Michael Feldbrügge, and Vera Göhre. 2017. 'The Plant-Dependent Life Cycle of Thecaphora Thlaspeos: A Smut Fungus Adapted to Brassicaceae'. *Molecular Plant-Microbe Interactions®* 30 (4): 271–82. https://doi.org/10.1094/MPMI-08-16-0164-R.

Gasciolli, Virginie, Allison C. Mallory, David P. Bartel, and Hervé Vaucheret. 2005. 'Partially Redundant Functions of Arabidopsis DICER-like Enzymes and a Role for DCL4 in Producing Trans-Acting siRNAs'. *Current Biology* 15 (16): 1494–1500. https://doi.org/10.1016/j.cub.2005.07.024.

Glawe, Dean A. 2008. 'The Powdery Mildews: A Review of the World's Most Familiar (Yet Poorly Known) Plant Pathogens'. *Annual Review of Phytopathology* 46 (Volume 46, 2008): 27–51. https://doi.org/10.1146/annurev.phyto.46.081407.104740.

Harvey, Jagger J. W., Mathew G. Lewsey, Kanu Patel, Jack Westwood, Susanne Heimstädt, John P. Carr, and David C. Baulcombe. 2011. 'An Antiviral Defense Role of AGO2 in Plants'. *PLOS ONE* 6 (1): e14639. https://doi.org/10.1371/journal.pone.0014639.

Jaubert, Marianne, Saikat Bhattacharjee, Alexandre F.S. Mello, Keith L. Perry, and Peter Moffett. 2011. 'ARGONAUTE2 Mediates RNA-Silencing Antiviral Defenses against Potato Virus X in Arabidopsis'. *Plant Physiology* 156 (3): 1556–64. https://doi.org/10.1104/pp.111.178012.

Landa, Blanca B., Maria Saponari, Oseias R. Feitosa-Junior, Annalisa Giampetruzzi, Filipe J. D. Vieira, Eliana Mor, and Silke Robatzek. 2022. 'Xylella Fastidiosa's Relationships: The Bacterium, the Host Plants, and the Plant Microbiome'. *New Phytologist* 234 (5): 1598–1605. https://doi.org/10.1111/nph.18089.

Liu, Wenxian, Hongmei Zeng, Zhipeng Liu, Xiufen Yang, Lihua Guo, and Dewen Qiu. 2014. 'Mutational Analysis of the *Verticillium Dahliae* Protein Elicitor PevD1 Identifies

Distinctive Regions Responsible for Hypersensitive Response and Systemic Acquired Resistance in Tobacco'. *Microbiological Research* 169 (5): 476–82. https://doi.org/10.1016/j.micres.2013.08.001.

Luneau, Julien S., Laurent D. Noël, Emmanuelle Lauber, and Alice Boulanger. 2022. 'A β-Glucuronidase (GUS) Based Bacterial Competition Assay to Assess Fine Differencesin Fitness during Plant Infection'. *Bio-Protocol* 12 (13): e3776. https://doi.org/10.21769/BioProtoc.3776.

Mallory, Allison C., Annika Hinze, Matthew R. Tucker, Nicolas Bouché, Virginie Gasciolli, Taline Elmayan, Dominique Lauressergues, Vincent Jauvion, Hervé Vaucheret, and Thomas Laux. 2009. 'Redundant and Specific Roles of the ARGONAUTE Proteins AGO1 and ZLL in Development and Small RNA-Directed Gene Silencing'. *PLOS Genetics* 5 (9): e1000646. https://doi.org/10.1371/journal.pgen.1000646.

Mitre, Letícia Kuster, Natália Sousa Teixeira-Silva, Katarzyna Rybak, Diogo Maciel Magalhães, Reinaldo Rodrigues de Souza-Neto, Silke Robatzek, Cyril Zipfel, and Alessandra Alves de Souza. 2021. 'The Arabidopsis Immune Receptor EFR Increases Resistance to the Bacterial Pathogens Xanthomonas and Xylella in Transgenic Sweet Orange'. *Plant Biotechnology Journal* 19 (7): 1294–96. https://doi.org/10.1111/pbi.13629.

Morel, Jean-Benoit, Christian Godon, Philippe Mourrain, Christophe Béclin, Stéphanie Boutet, Frank Feuerbach, Florence Proux, and Hervé Vaucheret. 2002. 'Fertile Hypomorphic ARGONAUTE (Ago1) Mutants Impaired in Post-Transcriptional Gene Silencing and Virus Resistance'. *The Plant Cell* 14 (3): 629–39. https://doi.org/10.1105/tpc.010358.

O'Connell, Richard J., and Ralph Panstruga. 2006. 'Tête à Tête inside a Plant Cell: Establishing Compatibility between Plants and Biotrophic Fungi and Oomycetes'. *The New Phytologist* 171 (4): 699–718. https://doi.org/10.1111/j.1469-8137.2006.01829.x.

Osborne, Rory, Laura Rehneke, Silke Lehmann, Jemma Roberts, Melina Altmann, Stefan Altmann, Yingqi Zhang, et al. 2023. 'Symbiont-Host Interactome Mapping Reveals Effector-Targeted Modulation of Hormone Networks and Activation of Growth Promotion'. *Nature Communications* 14 (1): 4065. https://doi.org/10.1038/s41467-023-39885-5.

Plücker, Lesley, Kristin Bösch, Lea Geißl, Philipp Hoffmann, and Vera Göhre. 2021. 'Genetic Manipulation of the Brassicaceae Smut Fungus Thecaphora Thlaspeos'. *Journal of Fungi* 7 (1): 38. https://doi.org/10.3390/jof7010038.

Qiang, Xiaoyu, Michael Weiss, Karl- Heinz Kogel, and Patrick Schäfer. 2011. 'Piriformospora Indica—a Mutualistic Basidiomycete with an Exceptionally Large Plant Host Range'. *Molecular Plant Pathology* 13 (5): 508–18. https://doi.org/10.1111/j.1364-3703.2011.00764.x.

Qu, Feng, Xiaohong Ye, and T. Jack Morris. 2008. 'Arabidopsis DRB4, AGO1, AGO7, and RDR6 Participate in a DCL4-Initiated Antiviral RNA Silencing Pathway Negatively Regulated by DCL1'. *Proceedings of the National Academy of Sciences of the United States of America* 105 (38): 14732–37. https://doi.org/10.1073/pnas.0805760105.

Roper, Caroline, Claudia Castro, and Brian Ingel. 2019. '*Xylella Fastidiosa*: Bacterial Parasitism with Hallmarks of Commensalism'. *Current Opinion in Plant Biology*, Biotic interactions, 50 (August):140–47. https://doi.org/10.1016/j.pbi.2019.05.005.

Singh, Jasleen, Vibhor Mishra, Feng Wang, Hsiao-Yun Huang, and Craig S. Pikaard. 2019. 'Reaction Mechanisms of Pol IV, RDR2, and DCL3 Drive RNA Channeling in the siRNA-Directed DNA Methylation Pathway'. *Molecular Cell* 75 (3): 576-589.e5. https://doi.org/10.1016/j.molcel.2019.07.008.

Spanu, Pietro D., James C. Abbott, Joelle Amselem, Timothy A. Burgis, Darren M. Soanes, Kurt Stüber, Emiel Ver Loren van Themaat, et al. 2010. 'Genome Expansion and Gene Loss in Powdery Mildew Fungi Reveal Tradeoffs in Extreme Parasitism'. *Science* 330 (6010): 1543–46. https://doi.org/10.1126/science.1194573.

Vaucheret, Hervé. 2006. 'Post-Transcriptional Small RNA Pathways in Plants: Mechanisms and Regulations'. *Genes & Development* 20 (7): 759–71. https://doi.org/10.1101/gad.1410506.

Xie, Zhixin, Lisa K. Johansen, Adam M. Gustafson, Kristin D. Kasschau, Andrew D. Lellis, Daniel Zilberman, Steven E. Jacobsen, and James C. Carrington. 2004. 'Genetic and Functional Diversification of Small RNA Pathways in Plants'. *PLoS Biology* 2 (5). https://doi.org/10.1371/journal.pbio.0020104.

Xin, Xiu-Fang, and Sheng Yang He. 2013. '*Pseudomonas Syringae* Pv. *Tomato* DC3000: A Model Pathogen for Probing Disease Susceptibility and Hormone Signaling in Plants'. *Annual Review of Phytopathology* 51 (1): 473–98. https://doi.org/10.1146/annurev-phyto-082712-102321.

Zhang, Rui, Ying Jing, Haiyang Zhang, Yahan Niu, Chang Liu, Jin Wang, Ke Zen, Chen-Yu Zhang, and Donghai Li. 2018. 'Comprehensive Evolutionary Analysis of the Major RNA-Induced Silencing Complex Members'. *Scientific Reports* 8 (1): 14189. https://doi.org/10.1038/s41598-018-32635-4.
